# Changes in microhabitat structure around amphibian breeding ponds in the northern Rocky Mountains following severe wildfire

**DOI:** 10.64898/2026.01.03.697496

**Authors:** Tristan Skretting, Ashley M. Smith, Jenny L. McCune, Julie A. Lee-Yaw

**Affiliations:** Department of Biological Sciences, University of Lethbridge, 4401 University Dr W, Lethbridge, Alberta, T1K 6T5, Canada; Department of Biology, University of Ottawa, 30 Marie-Curie, Ottawa, Ontario, K1N 9B4, Canada

**Keywords:** disturbance, BACI, Rocky Mountains, long-toed salamander, western toad, Columbia spotted frog, boreal chorus frog

## Abstract

Microhabitats promote diverse and resilient forests. Wildfire may alter critical microhabitat, yet studies directly quantifying these effects are limited. Here, we use a rare before-after-control-impact dataset to assess the impacts of severe wildfire on fine-scale gradients in microhabitat structure associated with amphibian breeding ponds in the Rocky Mountains. Using 462 photoquadrats sampled eight years before and two years after the 2017 Kenow wildfire in Waterton Lakes National Park, we quantified changes in 14 microhabitat features at three different distances (0, 3, and 10 m) away from burned and unburned ponds. Non-parametric multivariate analysis of variance suggested distance from pond edge was a significant predictor of microhabitat composition prior to the wildfire, with the proportion of soil (bare ground) decreasing and the proportion of shrubs and forbs increasing away from pond edges. A significant interaction between distance, burn status, and time was observed, suggesting that the relationship between microhabitat and distance depended on the joint effects of burn status and time. This result was driven by changes 10 m away from pond edges in the burn zone, where sites experienced an increase in the proportion of forbs, soil, gravel, and rocks, and the loss of moss cover following the fire. Microhabitat change was more pronounced at high elevation sites, although power to statistically test the effect of elevation was limited. Our study demonstrates heterogeneity in the impacts of wildfire on microhabitats along fine-scale gradients. We discuss implications of these findings for the management of amphibians using these sites.

## Introduction

The ecological importance of microhabitats is widely recognized. For small-bodied species, microhabitats provide refuge and resources, and buffer individuals against exposure to environmental extremes (De Frenne et al., 2021; Scheffers et al., 2014). Changes to microhabitats can have large effects on patterns of occupancy and abundance in some species (Ehlers Smith et al., 2017; Williams & Alexander, 2023; Zou et al., 2017), making it important to understand where and how global change impacts the microhabitats used by sensitive wildlife taxa (De Frenne et al., 2021, 2025; Ellis, 2018; Sallé et al., 2021).

Although wildfires are a natural part of the disturbance regime of many ecosystems, climate change coupled with a history of colonial fire suppression has led to an increase in the frequency of high-severity wildfires (Coogan et al., 2019; Liu et al., 2010). Severe wildfires can alter microhabitats in a variety of ways (Wolf et al., 2021). In the immediate years following a severe burn, the loss of canopy and vegetative cover is expected to increase soil temperatures (Ebel, 2012; Hossack et al., 2009). Severe wildfire can also alter soil properties (Agbeshie et al., 2022) and the movement of water (Guzmán-Rojo et al., 2024), and thus affect substrate moisture levels. Finally, post-fire changes in ground-level plant communities (Dickson-Hoyle et al., 2024) and larger ecosystem transitions (Davis et al., 2019), may fundamentally alter the availability of specific microhabitats. These changes in turn may have consequences for the distribution, movement, and performance of small-bodied animals following major wildfire events (Driscoll et al., 2012; Lees et al., 2022; Plavsic, 2014; Santos et al., 2014).

Amphibians require cool, moist, and protected sites to thrive and tend to have strong microhabitat associations (e.g. Farallo & Miles, 2016; McAlpine & Dilworth, 1989; Thorpe et al., 2018). As a result, amphibian populations are expected to be highly sensitive to microhabitat change or loss, including changes associated with wildfire (e.g. Bailey et al., 2025; Bury, 2004; Gade et al., 2019; Muñoz et al., 2019; Papp & Papp, 2000). Structural changes to microhabitats may be particularly impactful for amphibians. For example, woody debris offers refuge for individuals in the terrestrial environment and consumption of such features by wildfire may put amphibians at greater risk of desiccation and/or predation (reviewed by Pilliod et al., 2003). Likewise, patterns of aggregation on the landscape may change if the availability of such cover objects become limiting (e.g. Cummer & Painter, 2007), influencing the frequency of intra- and interspecific interactions. Other structural changes to microhabitats following wildfire may impact other aspects of amphibian physiology and ecology. For instance, the loss of vegetative cover may increase the amount of solar radiation and the temperatures that amphibians are exposed to, while the loss of duff and leaf litter may influence foraging and the movement of individuals (reviewed by Pilliod et al., 2003; Brown, 2013). Assessing changes in microhabitat composition and structure following severe wildfire events is thus relevant to understanding the effects of these events on amphibian populations.

In this study, we use photos of quadrats placed around amphibian breeding ponds before and after a severe wildfire event in western Canada to characterize and compare changes in microhabitat composition in burned and unburned areas. Fine-scale moisture, temperature, and light gradients are expected to cause microhabitats to vary with distance from the edge of wetlands (e.g. Torralvo et al., 2022) and these gradients may influence the dispersion of different amphibian life stages and species around breeding ponds (e.g. Rittenhouse & Semlitsch, 2007). Thus, we first asked about the relationship between microhabitat composition and distance from pond edges and whether this relationship was altered by wildfire. We then explored the amount of change in microhabitat composition within sites. The relative importance of different microhabitat features for amphibians may vary across elevation (e.g. Farallo et al., 2018), therefore we also asked whether the amount of change observed varied across elevation. Our study addresses timely questions as to the impacts of severe wildfire on microhabitats around forest pond margins while simultaneously providing quantitative information about amphibian microhabitats in the southern Canadian Rocky Mountains—a region for which such information is sparse.

## Methods

### Study region and wildfire

Waterton Lakes National Park (WLNP) or *Paahtómahksikimi* (Blackfoot) is a 505 km^2^ federally protected area in southwestern Alberta, Canada (N 49° 03’ 0.288, W 113° 54’ 40.083”). Located along the Continental Divide in one of the narrowest parts of the Rocky Mountains, WLNP includes four ecoregions: foothills parkland, montane, subalpine, and alpine. These ecoregions differ with respect to the dominant vegetation, with fescue grasses and aspen being characteristic of the foothills parkland, coniferous forests (in particular, lodgepole pine, *Pinus contorta*) dominating montane and subalpine areas, and low-growing vegetation characterizing alpine areas (Strong & Leggat, 1992). The park experiences moist, relatively mild winters and short, cool summers, with mean annual air temperatures of 2.4 °C and annual precipitation of ∼1072 mm (Parks Canada, 2025a). Precipitation varies in the park, with western parts of the park at the continental divide receiving more annual precipitation (1520 mm) than eastern parts of the park (76 mm; Parks Canada, 2025a). Six amphibian species are found in WLNP including two salamanders (long-toed salamander: *Ambystoma macrodactylum*; western tiger salamander: *Ambystoma mavortium mavortium*), one toad (western toad: *Anaxyrus boreas*), and three frogs (Columbia spotted frog: *Rana luteiventris*; northern leopard frog: *Lithobates pipiens*; boreal chorus frog: *Pseudacris maculata*). In addition to these species, the park is home to over 1000 species of vascular plants, ∼ 60 species of mammals, ∼250 species of birds, 24 species of fish, and four species of reptiles (Parks Canada, 2025b), making it one of the most biodiverse regions in Alberta.

Forested areas in WLNP historically had fire return intervals of 26 to 85 years (depending on ecoregion: Rogeau, 2016) and experienced both mixed-severity and stand-replacing fires (Barrett, 1996); earlier fire history in the foothills parkland ecosystem also included intentional fires set by Indigenous peoples as discussed in (Eisenberg et al., 2019). However, colonial fire suppression practices led to late successional stands and forest encroachment throughout much of the park (Stockdale et al., 2019; Trant et al., 2020), with the last major fire prior to this study (1998 Sofa Mountain fire, 1521 hectares) burning stands that were over 130 years old (O’Connor, 2015). In late summer of 2017, the record-breaking Kenow Wildfire burned 19,303 hectares (∼50% of vegetated areas) of WLNP. This was a fast-moving fire (most of the area impacted was burned over the course of a single night) that resulted in an unusually uniform and severe burn, with 75% of the burned area classified as extreme severity (Greenaway et al., 2018). The wildfire was followed by several exceptionally dry years (Aspinall et al., 2025), likely increasing the importance of microhabitat refuges for amphibians in the burn footprint.

### Data collection

We had visited 20 amphibian breeding ponds in WLNP in July 2009 (Fig. 1a), eight years prior to the Kenow wildfire, as part of other research activities focused on the long-toed salamander. Although breeding ponds were selected based on the distribution of this species, most of these sites are used by other members of the amphibian community in southwestern Alberta (Table A1; although both tiger salamanders and the recently, reintroduced northern leopard frog occupy a limited number of sites in the park). Sites ranged in elevation from 1273 to 1973 m and were primarily located in the montane (N = 12) and foothills parkland (N = 5) ecoregions (Table A1). Selected breeding ponds were small (largest pond was a maximum of ∼ 200 m from one side to the other) and easily circumnavigated within one hour. At each site, we established four, ten meter transects radiating away from the focal pond on different sides of the pond (Fig. 1b). Along each transect, 1 m^2^ quadrat frames were placed at 0, 3, and 10 m away from the pond edge (Fig. 1b). A digital photo of each quadrat was taken from breast height, approximately parallel to the slope of the ground (to the extent possible). By chance, the Kenow wildfire burned nine of these survey sites, providing an opportunity to explore general changes to microhabitat structure around these sites based on data collected from these photos and to compare with unburned sites. In July 2020, three-years post-fire, we revisited all 20 sites and collected post-fire quadrat photos. Although waypoints at each location along the original 2009 transects were not recorded, we had GPS coordinates taken from the 0 m quadrat of the first 2009 transect at each site, as well as photos capturing a view of the wetland taken from beside each of the 0 m quadrats from the 2009 surveys. Using these coordinates and landmarks in the site photos (e.g. ridge lines, rocky outcrops, etc.) to guide transect starting locations, we positioned the 2020 transects as closely as possible to the 2009 transects to minimize the effects of choice of transect location when comparing microhabitat composition across time at each site; though we note that transects were not sufficiently matched to enable paired analysis at the transect or quadrat level.

**Figure 1.**
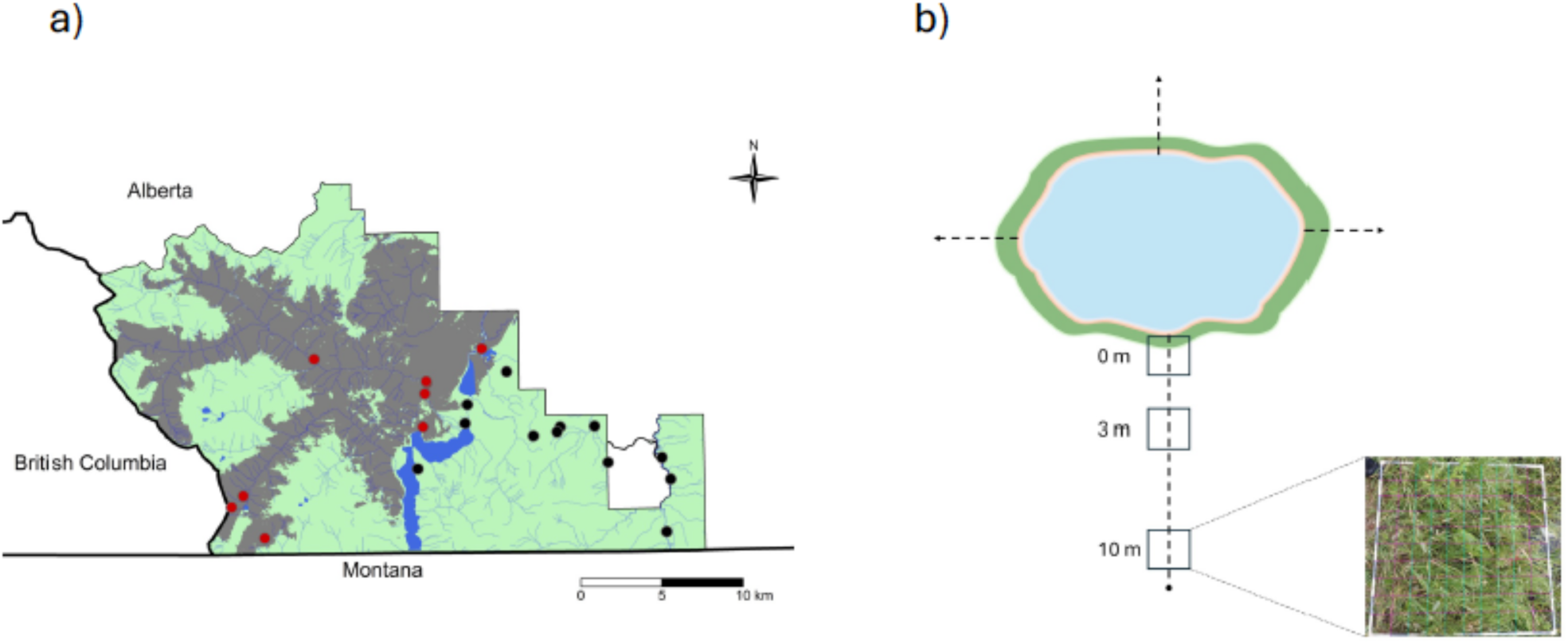
**a)** Distribution of the 20 amphibian breeding ponds included in microhabitat surveys in and adjacent to Waterton Lakes National Park (coloured area), with respect to the burn footprint of the 2017 Kenow wildfire (grey shading). Colours represent the burn status (red: 9 burned; black: 12 unburned) of each site. **b)** Schematic of the survey design at each site. Four transects were placed on different sides of each pond (single transect depicted here). Photos of 1 m^2^ quadrats (example in inset) were taken along each transect at 0 m, 3 m, and 10 m away from the pond edge, allowing for digital overlay of a grid used to get counts of 14 different microhabitat features within each quadrat (see Methods and Table 1).

**Table 1.** Features used to characterize microhabitat around wetlands in Waterton Lakes National Park, Alberta, Canada.

| Microhabitat Feature | Description | Rationale and focal species<sup>±</sup> examples<sup>†</sup> |
| --- | --- | --- |
| Soil | Exposed mineral soil | Open substrates influence movement and habitat selection (e.g. NLFR: (Blomquist & Hunter, 2009); LTSA: (Lee-Yaw et al., 2015) |
| Gravel | Small pebbles and rocks (<10 cm in diameter) | Open substrates influence movement decisions (e.g. LTSA: Lee-Yaw et al., 2015) and may influence burrowing (e.g. LTSA: (Sheppard, 1977)) |
| Rock | Rock >10 cm in diameter, large enough to shelter adult salamanders (~7 cm) | Provides cover (LTSA: (Jaeger, 1980); WETO: (Bull, 2006) |
| Litter | Accumulated organic material from decaying leaves, grass, bark, or needles | Influences movement decisions and habitat selection (e.g. NLFR: Blomquist and Hunter et al. 2009; LTSA: Jaeger 1980; Lee-Yaw et al. 2015); Associated with abundance (LTSA: (Graham, 1997) |
| Snag | Upright decaying wood or standing stump | Provides cover and overwintering sites (LTSA: (Atkinson-Adams et al., 2018) |
| Coarse woody debris (CWD) | Decaying downed wood >10 cm in diameter, large enough to shelter the adult salamanders (~7 cm) | Impacts on movement decisions (e.g. LTSA: (Hoza & Bakker, 2023) and habitat selection (e.g. NLFR: Blomquist and Hunter et al. 2009; WETO: (Browne & Paszkowski, 2018) |
| Moss | Non-vascular flowerless plants in taxonomic division *Bryophyta* (e.g. *Ambylstegium varium*) | Influences movement and habitat selection (e.g. LTSA: Lee-Yaw et al. 2015); Associated with abundance (e.g. WETO: (Browne et al., 2009) |
| Grass | Graminoids in family *Poaceae* (e.g. *Festuca scrabella*) | Influences movement and habitat selection (e.g. LTSA: Lee-Yaw et al. 2015); associated with abundance (e.g. BCRF: (Ouellet et al., 2009) |
| Sedge | Grass-like families *Cyperaceae* and *Juncaceae* (e.g. *Carex utriculate*) | Associated with abundance (e.g. BCRF: Ouellet et al. 2009) and distribution (e.g. CSFR: (Shive et al., 2010) |
| Forb | Non-woody, non-graminoid flowering plants (e.g. *Arnica cordifolia*) | Herbaceous cover influences distribution (e.g. BCRF: Constible et al. 2001) |
| *Equisetum* spp. | Plants in genus *Equisetum* (e.g. *Equisetum arvense*) | Herbaceous cover influences distribution (e.g. BCRF: (Constible et al., 2001) |
| Shrub | Woody plants, usually multi-stemmed and less than 10 m when mature (e.g. *Cornus sericea*) | Influences distribution (BCFR: Constible et al. 2001) and habitat selection (e.g. WETO: Browne and Paszkowski 2018) and abundance (BCFR, WETO: Browne et al. 2009) |
| Tree:<br>Deciduous | Deciduous tree species (e.g. *Populus tremuloides*) | Influences abundance (e.g. WETO, BCFR: Browne et al.); Canopy cover influences habitat selection (NLFR: Blomquist and Hunter et al. 2009) and abundance (BCFR, WETO: Browne et al. 2009) |
| Tree:<br>Coniferous | Coniferous tree species (e.g. *Pseudotsuga menziesii*) | Influences abundance (e.g. WETO, BCRF: Browne et al.); Canopy cover influences habitat selection (NLFR: Blomquist and Hunter et al. 2009) and abundance (BCFR, WETO: Browne et al. 2009) |

<sup>±</sup> BCFR = boreal chorus frog (*Pseudacris maculata*); CSFR = Columbia spotted frog (*Rana luteiventris*); LTSA = long-toed salamander (*Ambystoma macrodactylum*); NLRf = northern leopard frog (*Lithobates pipiens*); WETO = western toad (*Anaxyrus boreas*)
<sup>†</sup>Associations indicated may be positive or negative depending on species.

Each site had a total of 24 photoquadrats (four transects, three distance classes, two time-points), with the exception of two sites for which one side of the wetland was inaccessible (only 18 quadrats sampled for these site) and two sites, each with a single missing or blurry quadrat photo that could not be used (only 23 quadrats sampled for these sites). Photoquadrats were imported into Adobe Photoshop (version 21.2.3). Quadrat frames had physical markings at 10 cm intervals along each side that were used to digitally superimpose a 10 x 10 cm grid over each quadrat (Fig. 1b). We classified each of the 81 internal points formed by the intersecting grid lines into one of 14 microhabitat features (Table 1; see Booth et al., 2006; Degrassi, 2018; Howes & Lougheed, 2004 for similar methods). Focal microhabitat features fall into the broader categories of bare ground, dead organic or inorganic cover, moss, herbaceous vegetation, and woody-stemmed vegetation, all of which have previously been shown to impact occupancy, abundance, and movement in temperate amphibians, including the species found in WLNP (Table 1). Quadrats where more than 20% of counts corresponded with submerged terrain or fell on roads (i.e. “nuisance” categories) were removed prior to analysis (three quadrats from two sites). A total of 462 quadrats were retained in our analysis (Table A1).

### General analytical approach

The nature of our dataset (counts of features within quadrats) lends itself to multivariate analysis and our general approach was as follows. To prepare the data for analyses, we first converted counts of each microhabitat feature to proportions, ignoring remaining “nuisance” counts when calculating the total number of counts in each quadrat. Because we are focused on compositional differences among samples with many zeros, we transformed the data using a Hellinger transformation (Legendre & Gallagher, 2001). We visualized microhabitat composition in multivariate space using non-metric multidimensional scaling (NMDS) based on Bray-Curtis dissimilarities of the Hellinger-transformed data for all quadrats and fit environmental vectors onto the resulting ordination using the “metaMDS” and “envfit” functions respectively in the *vegan* package (version 2.7-1; Oksanen et al., 2025) in R (version 4.4.3; R Core Team 2025). Our questions involved testing the effects of categorical predictors on multivariate microhabitat composition (e.g. distance class, burn status, etc.). To do so, we used permutational multivariate analysis of variance (PERMANOVA). PERMANOVA specifically tests for differences in the centroids and/or dispersion of specified groups in multivariate space (Anderson, 2017). Significant PERMANOVA results were followed by tests for homogeneity of dispersion, as well as post-hoc tests (pairwise ANOVAs) to identify specific groups that differ. PERMANOVAs, dispersion tests, and post-hoc tests were based on Bray-Curtis dissimilarities of the Hellinger-transformed data and run in PRIMER-e (Anderson et al., 2008). Significant PERMANOVAs were also followed by an indicator analysis to identify the microhabitat features most strongly associated with groups (Dufrêne & Legendre, 1997). Indicator analyses were run using the “multipatt” function in the *indicspecies* package (version 1.8.0; De Cáceres & Legendre, 2009) with the association function set to “r.g”. We set the number of permutations to 5000 in all permutation-based analyses.

### Relationship between distance to pond edge and microhabitat composition

We first asked whether microhabitat composition varied among the three distance classes prior to the fire (i.e. baseline conditions). To address this question, we used PERMANOVA to test the effects of distance class on multivariate dissimilarity among quadrats in the 2009 dataset. To account for non-independence associated with quadrats within sites, we included site as a random effect in this model, allowing quadrats to be shuffled across distance classes within site to test the significance of distance class while preserving the assignment of quadrat to site.

The removal of woody vegetation during the wildfire might be expected to homogenize microhabitat composition across distance classes within in the burn zone, at least in the short-term. Such an effect would be evident from both the loss of any relationship between microhabitat composition and distance class, as well as a reduction in multivariate dispersion across sites and distances in the burn zone after the fire. To assess whether the relationship between microhabitat composition and distance class in the burn zone changed following the fire, we used PERMANOVA to test the three-way interaction between distance class, time, and burn status. To simplify the analysis, we aggregated counts from quadrats from the same distance class and time period within sites prior to calculating proportions and transforming the data (hereafter referred to as the “aggregated dataset”). Site was included as a random effect in this analysis to account for non-independence associated with repeated measures of the same site over time. Post-hoc tests were used to explore meaningful comparisons (e.g. significant differences between pre- and post-fire composition within burn status and/or differences between burned and unburned sites within each time period) and an indicator analysis was used to look for microhabitat features significantly associated with different distance x burn status x time groups. For the latter, a custom hierarchical permutation scheme was used to account for non-independence associated with site and repeated measures within site. Specifically, sites were shuffled as intact units, time period was shuffled within site, and distance class shuffled within time period within site. Significance was assessed by comparing observed association statistics from *multipatt* to a null distribution of values based on the permuted dataset. We also tested for differences in dispersion among groups, asking whether multivariate dispersion was lower for post-fire samples than for pre-fire samples at the same sites (i.e. expected if there has been homogenization over time), and especially lower for post-fire samples in burned areas relative to unburned areas (i.e. expected if the wildfire and not just time is the cause of any homogenization).

### Differences in the amount of change among sites

We next tested potential drivers of the amount of change in microhabitat composition over time. We calculated the Bray-Curtis dissimilarity between the 2009 and 2020 surveys at each distance class and used the dissimilarity between time points at a given site as an estimate of the amount of change in microhabitat composition at that distance class at that site (e.g. Lloren & McCune, 2024). Dissimilarities range from 0 (no discernable change) to 1, with larger values indicating a greater amount of change. We used Spearman Rank correlation to assess whether there were associations between the rank amount of change at one distance class and the rank amount of change at other distance classes within sites. For each distance class, we used generalized linear models with a beta distribution in the *betareg* package in R to assess the effects of burn status, elevation, and the interaction between burn status and elevation on the amount of change within sites. Given baseline differences in microhabitat composition among the different distance classes (see Section 3.1), we ran a separate analysis for each distance class, using the aggregated dataset to simplify the models. Elevation was centered to better meet model assumptions. NMDS plots based on the aggregated data and with the associated environmental vectors based on an envfit analysis for each distance class were generated to facilitate interpretation of results. Small sample sizes and lack of variation across levels of other predictors of interest (e.g. ecoregion and burn severity) precluded exploration of other ecological covariates that may moderate the effects of wildfire on microhabitats.

## Results

### Relationship between distance to pond edge and microhabitat composition

We first characterized baseline microhabitat structure and explored fine-scale gradients in microhabitat composition away from pond edges before the wildfire. PERMANOVA based on the 2009 data revealed significant differences in microhabitat composition at different distances away from pond edges (F_2, 57_ = 2.9441, p = 0.006; Table A2). A test of differences in multivariate dispersion suggested the significant PERMANOVA result was largely driven by differences in the mean (centroid) of groups rather than differences in dispersion (Table A3). Post-hoc tests point to differences in pre-fire microhabitat composition between the 0 and 10 m distance class (Table A4). Results from an indicator analysis suggest that quadrats adjacent to pond edges tended to have a greater proportion of sedge relative to quadrats 3 and 10 m away from the shoreline prior to the fire, whereas quadrats 10 m away from the shoreline had more deciduous trees (Table A5), though the latter made up a relatively small proportion of counts across the entire dataset (Fig. A1). There was more bare soil immediately at, and 3 m away, from the shoreline than 10 m away from the shoreline prior to the fire, and shrubs and forbs were more prominent at 3 and 10 m away from the shoreline than at the shoreline (Table A5; Fig. A1).

We next asked whether the relationship between microhabitat composition and distance changed following the wildfire and whether these changes depended on whether sites were burned. Consistent with an effect of the fire on the relationship between microhabitat composition and distance from pond edge, we found a significant three-way interaction in the PERMANOVA testing the joint effects of distance class, time, and burn status on microhabitat composition (Fig. 2, Table 2). Post-hoc tests revealed that this result was largely driven by changes in microhabitat composition at the 10 m distance class in the burn zone (Table A6). Visualization of mean change for each distance class in burned and unburned areas along the different axes resulting from the NMDS ordination of all quadrats suggest the greatest shift for the 10 m distance class was along axis 3 (Fig. 3). In particular, quadrats 10 m away from the pond edge in the burn zone showed decreases along axis 3 (Fig. 3), corresponding to shifts towards more forbs (Table A7; Fig. A1). A corresponding indicator analysis further suggests that mosses/lichen were significantly associated with all distance classes in the burn zone before the fire, but only with the 0 and 3 m distance classes in the burn zone after the fire (association coefficient = 0.373, p = 0.013; see also Fig. A1). Differences in multivariate dispersions among distance-burn status-time period groups were not significant (PERMDISP: F_11,108_ = 1.2871, p = 0.3975, nperm = 5000; Fig. A1).

**Figure 2.**
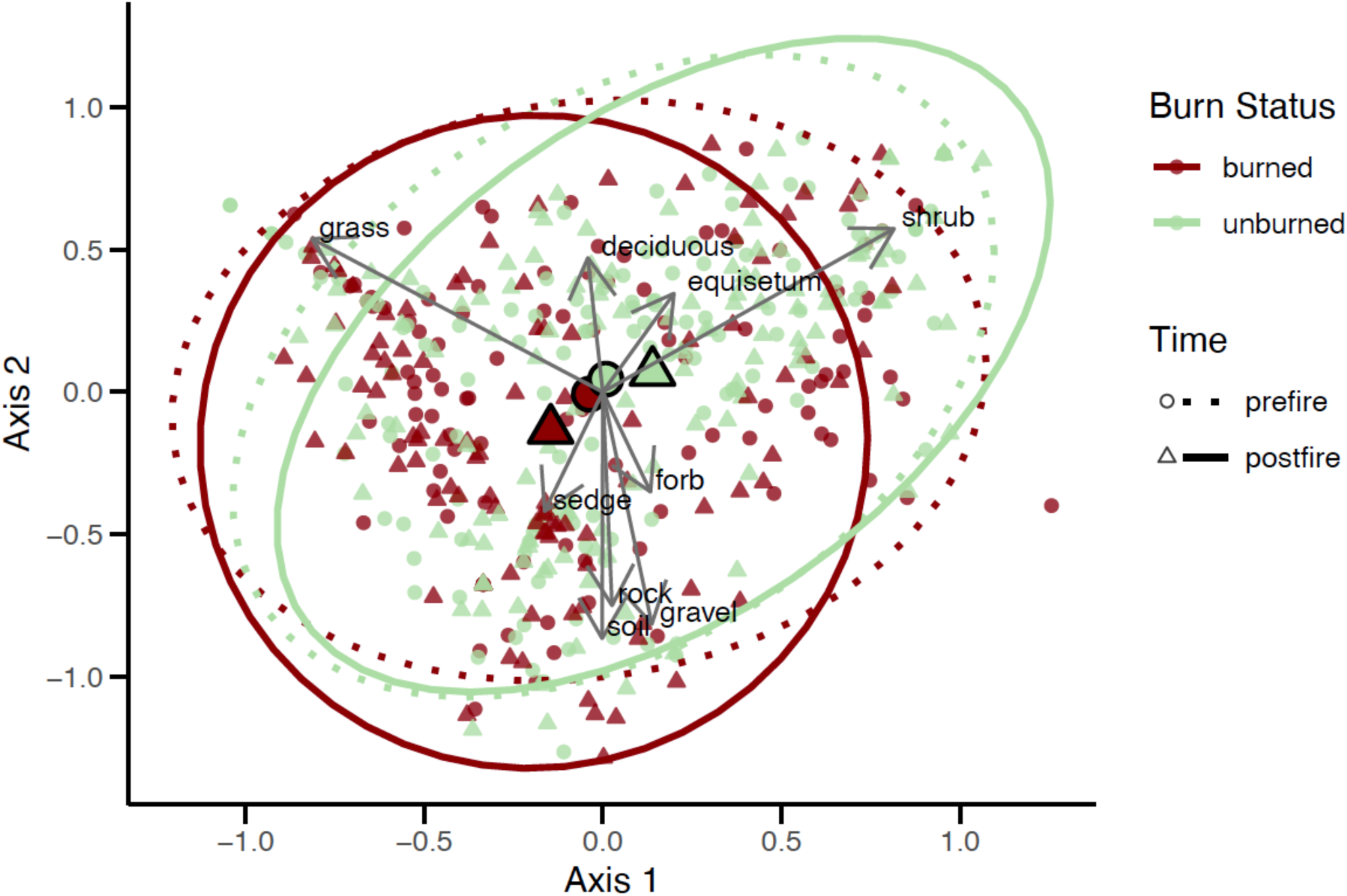
Non-metric multidimensional scaling (NMDS) ordination of microhabitat composition (NMDS based on Bray-Curtis dissimilarities; k = 3, stress = 0.125) among 1 m^2^ quadrats collected around amphibian breeding ponds in Waterton Lakes National Park before (circles) and after (triangles) the Kenow wildfire in burned (red) and unburned (green) parts of the park. For illustrative purposes, 95% confidence ellipses around burn status x time groups are shown, with group centroids indicated by the larger-sized symbols. Environmental vectors significantly associated with the first two axes are shown.

**Figure 3.**
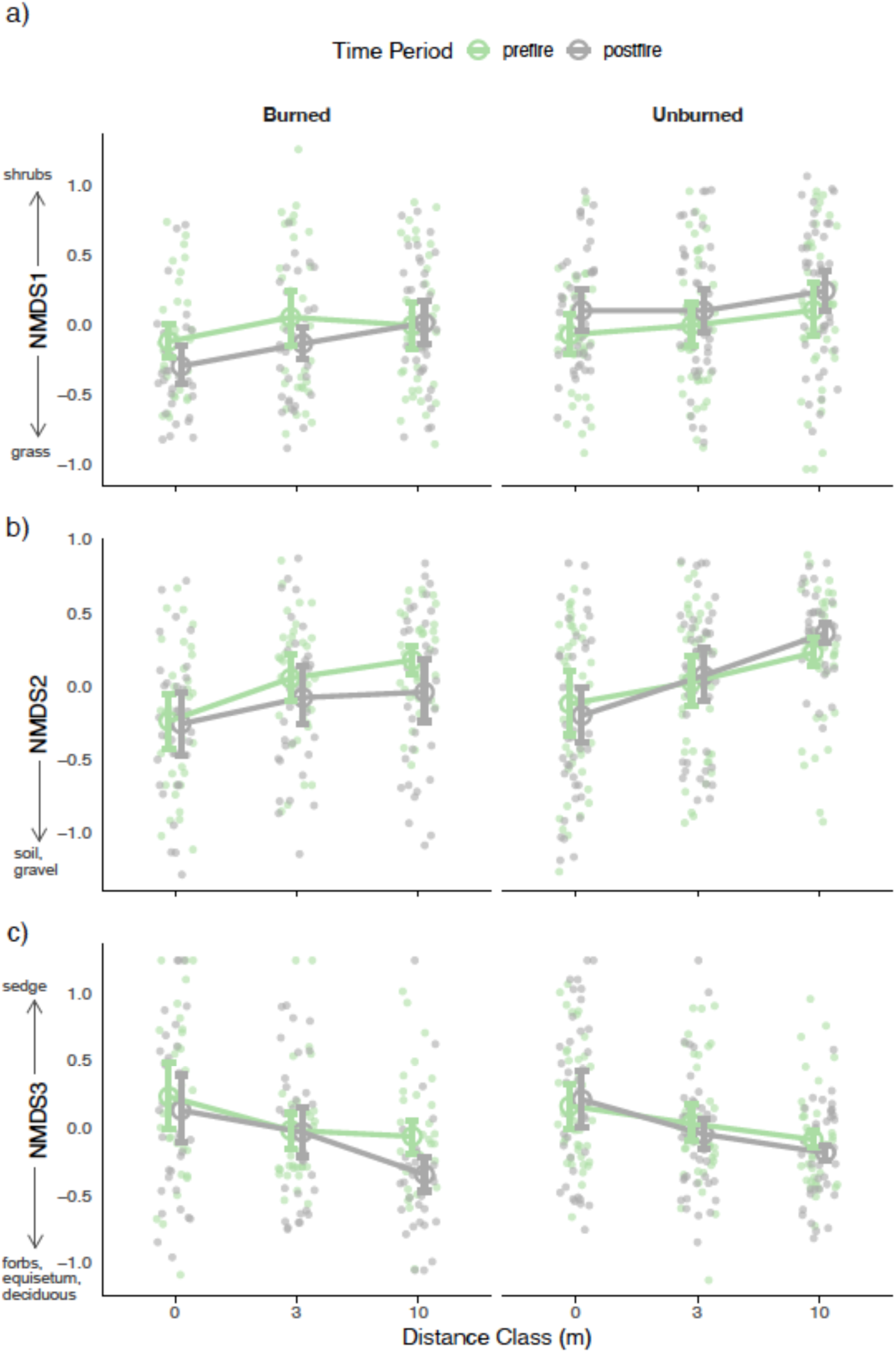
Relationship between microhabitat composition and distance away from pond edge (measured at 0, 3, and 10 m) before (green) and after (grey) the 2017 Kenow wildfire at nine burned (left panels) and eleven unburned (right panels) amphibian breeding sites. Each plot shows a different axis from a non-metric multidimensional scaling analysis summarizing the variation in the dataset (k = 3; stress = 0.125), with arrows along the y axes indicating environment vectors significantly and strongly associated with that axis (Table A7). Points represent values for individual 1 m^2^ quadrats. Error bars are 95% confidence intervals based on a bootstrapping scheme in which sites were resampled with replacement and mean values per distance class and time period were calculated per sampled site before averaging values across sampled sites within each distance-time-burn status group.

**Table 2.** Results from a PERMANOVA testing the effects of distance class (0, 3, or 10 m), time (pre- or post-fire), and burn status (burned or unburned) on differences in relative microhabitat composition at 20 amphibian breeding sites in southwestern Alberta.

| **Response** | **Predictor** | **df** | **SS** | **R<sup>2</sup>** | **F** | **p-value\*** |
| --- | --- | --- | --- | --- | --- | --- |
| Dissimilarity in relative microhabitat composition | Distance class | 2 | 13980 | 6990.1 | 11.518 | 0.00020 |
|  | Burn status | 1 | 3410.1 | 3410.1 | 0.94211 | 0.44630 |
|  | **Time** | **1** | **2230.2** | **2230.2** | **2.6361** | **0.04400** |
|  | **Site (Burn Status)** | **18** | **65154** | **3619.7** | **10.356** | **0.00020** |
|  | Distance class x Burn status | 2 | 656.96 | 328.48 | 0.54125 | 0.83060 |
|  | Distance class x Time | 2 | 326.58 | 163.29 | 0.46717 | 0.81720 |
|  | Burn status x Time | 1 | 2141.2 | 2141.2 | 2.5309 | 0.05640 |
|  | **Distance class x Site (Burn status)** | **36** | **21848** | **606.89** | **1.7363** | **0.00040** |
|  | **Time x Site (Burn status)** | **18** | **15229** | **846.04** | **2.4205** | **0.00020** |
|  | **Distance class x Burn status x Time** | **2** | **1899.7** | **949.85** | **2.7175** | **0.01320** |
|  | Residual | 36 | 12583 | 349.53 |  |  |
|  | Total | 119 | 1.39000 |  |  |  |

\*Based on 5000 permutations

### Magnitude of microhabitat change within sites

The magnitude of change in microhabitat composition (i.e. dissimilarity between time points in multivariate space) at adjacent distance classes was correlated (Spearman’s rho 0 to 3 m: 0.574, p = 0.009; Spearman’s rho 3 to 10 m: 1, p = 0.000006) but the magnitude of change at 0 and 10 m was not correlated (Spearman’s rho: 0, p = 0.841). Dissimilarity within sites across time at the shoreline (0 m) ranged from 0.117 to 0.585 (median = 0.295). Site shorelines did not tend to change in a consistent direction (Fig. 4a) and none of our predictors of interest were significantly associated with the amount of change within sites at this distance class (Table 3; Fig. 5a). For the 3 m distance class, dissimilarity between temporal samples from the same site ranged from 0.175 to 0.637 (median = 0.265). As per the 0 m distance class, sites did not change in a consistent fashion at 3 m away from pond edges (Fig. 4b) and none of our predictors were significantly associated with the amount of change at 3 m (Table 3; Fig. 5b). For the 10 m distance class, dissimilarity between temporal samples from the same site ranged from 0.120 to 0.624 (median = 0.314). Sites in the burn zone showed decreases along axis 2 of a NMDS plot based on this distance class (Fig. 4c), corresponding to increases in rocks, soil, and forbs (Table A8; see also Fig. A1). Our full model explained a substantial amount of the variation in change within sites for this distance class (Pseudo R2 = 0.388; Table 3). In this model, the interaction between burn status and elevation was marginally significant and the main effect of elevation was significant (Table 3). In general, the amount of change in microhabitat composition increased with elevation, and this appeared to be driven by three high elevation sites (sites 104, 105, and 106) in the burn zone (Fig. 5c). These sites had high leverage in all models and results for the 3 and 10 m distance class were sensitive to the sequential removal of these sites (Table A9). Of these sites, site 105 is particularly noteworthy as it demonstrated either the highest or second highest amount of change at all distance classes (Fig. 4).

**Figure 4.**
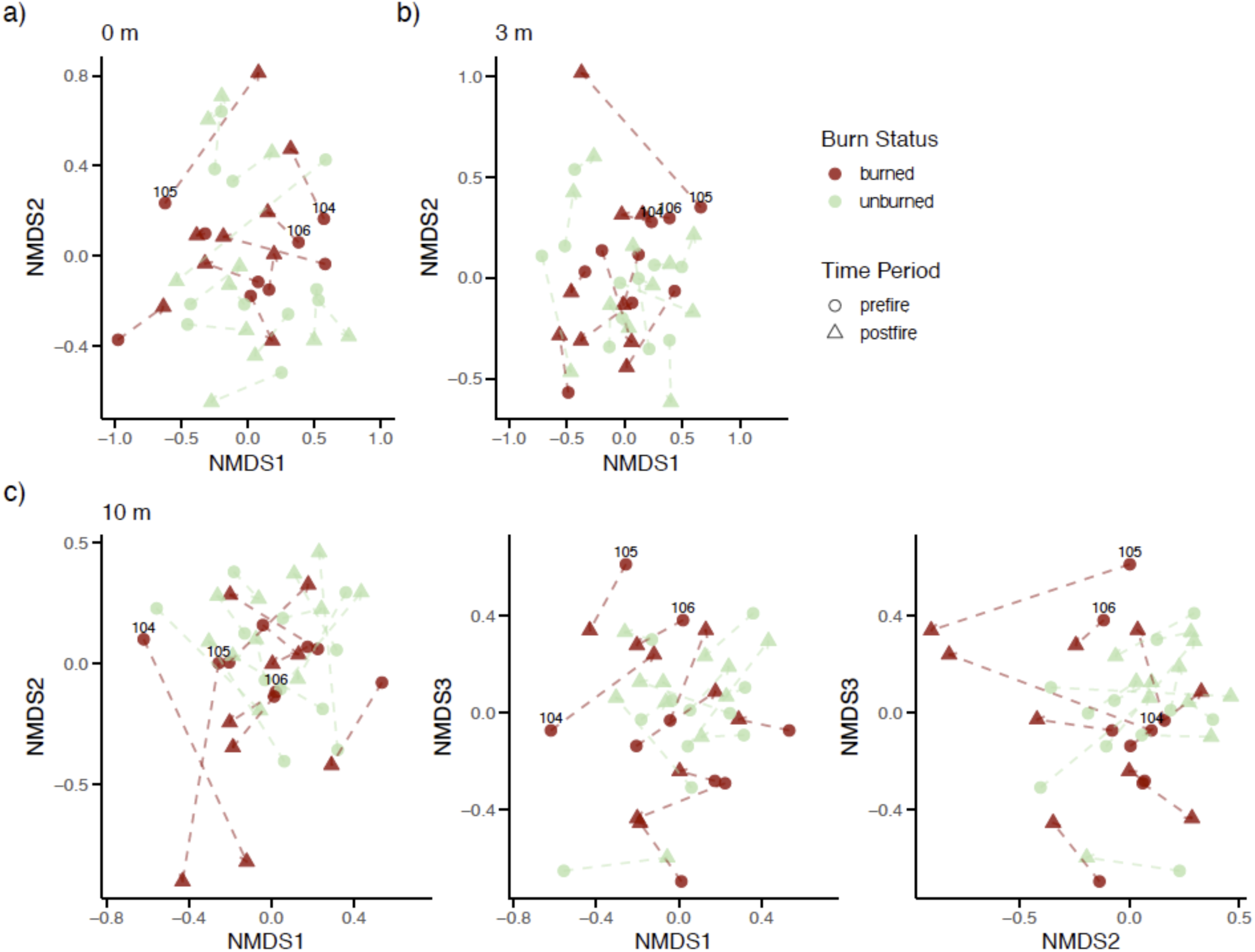
Non-metric multidimensional scaling (NMDS) plots showing shifts in microhabitat composition at a) 0 m, b) 3 m, and c) 10 m away around burned (red) and unburned (green) amphibian breeding ponds following a major wildfire in southwestern Alberta. Dashed lines connect each site before (circles) and after (triangles) the wildfire. NMDS plots are based on Bray-Curtis dissimilarities and were fit with k = 2 for the 0 m and 3 m distance classes and with k = 3 for the 10 m distance class. The three highest elevation sites in the burn zone are labeled.

**Table 3.** Results from linear models (beta regression) testing change in proportional microhabitat composition (pairwise Bray-Curtis dissimilarity) around amphibian breeding ponds in southwestern Alberta with respect to burn status and elevation. Significant predictors are bolded.

| **Response** | **Predictor** | **Coefficient\*** | **SE** | **Z** | **p-value\*\*** |
| --- | --- | --- | --- | --- | --- |
| Change in microhabitat composition 0 m away from pond edges<br>*Pseudo R-squared: 0.121* | Intercept | -0.871 | 0.182 | -4.78 |  |
|  | Burn status | -0.0258 | 0.248 | -0.10 | 0.373 |
|  | Elevation | 0.000948 | 0.000701 | 1.35 | 0.293 |
|  | Burn status x Elevation | -0.00224 | 0.00161 | -1.38 | 0.167 |
| Change in microhabitat composition 3 m away from pond edges<br>*Pseudo R-squared: 0.196* | Intercept | -0.620 | 0.147 | -4.225 |  |
|  | Burn status | -0.341 | 0.207 | -1.65 | 0.179 |
|  | Elevation | 0.000605 | 0.000577 | 1.048 | 0.430 |
|  | Burn status x Elevation | -0.00159 | 0.00138 | -1.15 | 0.247 |
| Change in microhabitat composition 10 m away from pond edges<br>*Pseudo R-squared: 0.388* | Intercept | -0.802 | 0.166 | -4.83 |  |
|  | Burn status | -0.0509 | 0.224 | -0.23 | 0.147 |
|  | **Elevation** | 0.00230 | 0.000633 | 3.63 | **0.00558** |
|  | Burn status x Elevation | -0.00292 | 0.00145 | -2.011 | 0.0506 |

\* Coefficients for burn status represent comparison to the reference category “unburned”.
\*\* p-values reflect marginal effects of predictor and were obtained from likelihood ratio tests comparing model with and without the predictor; to test each main effects, both the main effect and the interaction term were removed in the reduced model.

**Figure 5.**
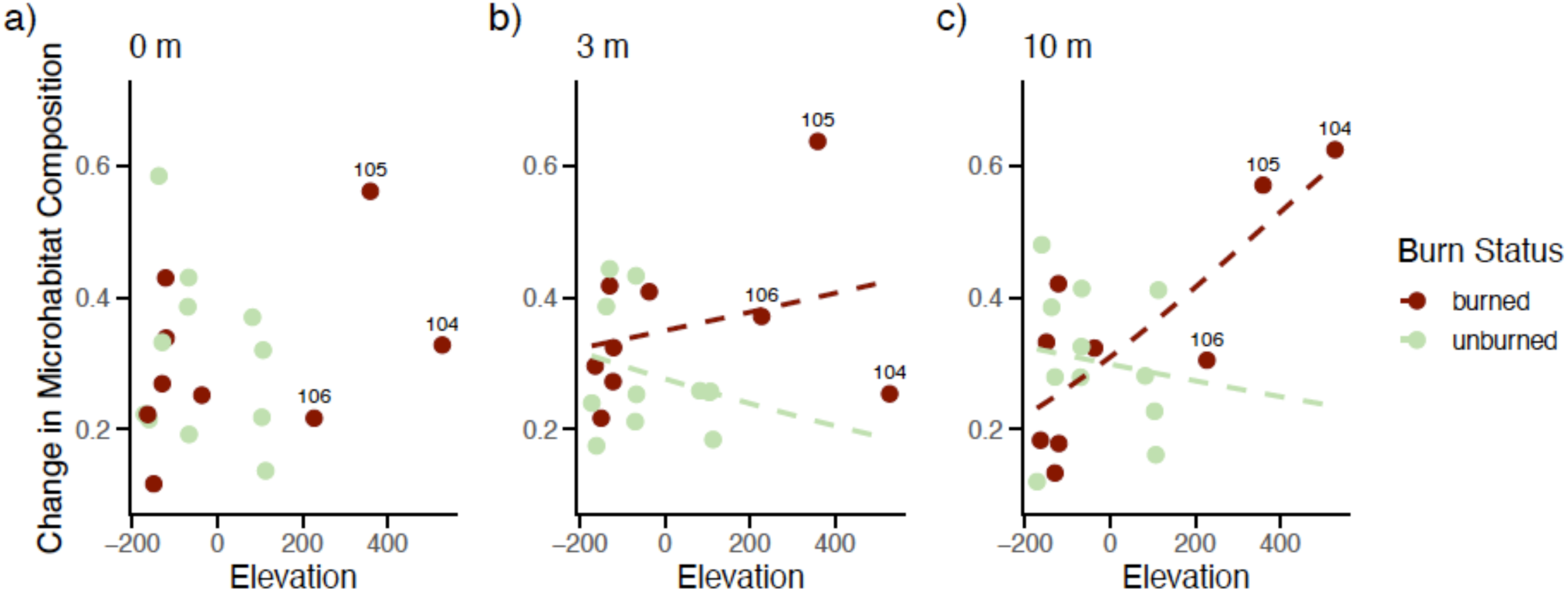
Change in multivariate microhabitat composition around amphibian breeding ponds at a) 0 m, b) 3 m, and c) 10 m away from the pond’s edge in relation to burn status and elevation (m, centered). Dashed lines in b) and c) show the interaction between burn status and elevation, the significance of which was sensitive to three high elevation sites in the burn zone (labeled in each panel; see Tables 4 and S7).

## Discussion

Understanding how microhabitats are altered by changing disturbance regimes is critical to our ability to predict and mitigate the effects of climate change on small-bodied taxa. Yet, studies quantifying the effects of large disturbance events on microhabitats are limited owing to the difficulty predicting these events. In this study, we took advantage of a rare opportunity to use a natural, before-after-control-impact design to examine the effects of a severe wildfire event on microhabitat structure around amphibian breeding ponds. Our results suggest that the Kenow wildfire had a limited effect on the composition of microhabitats immediately adjacent to breeding ponds in Waterton Lakes National Park. At the same time, we detected a wildfire effect as close as 10 m away from pond edges, suggesting that any buffering effects of wetlands in the park were restricted to very small spatial scales. Sites at high elevations also showed signs of being more impacted than sites at lower elevations. We discuss the potential implications of these results for amphibians in the park.

### Baseline microhabitat composition and changes following the wildfire

We first established a baseline understanding of microhabitat structure and composition around amphibian breeding ponds in Waterton Lakes National Park. We observed fine-scale gradients in microhabitat structure moving away from pond edges in the park before the Kenow wildfire, with near-shore quadrats (0 and 3 m) being characterized by more bare ground than quadrats further away (10 m) from pond edges. Although there was considerable variation among quadrats (see Fig. 2), in general vegetation transitioned away from more sedges around pond edges, to increasing shrubs and forbs by 3 m, to more deciduous trees by 10 m. Thus, prior to the wildfire, gradients in vegetation with distance from pond edges occurred over very fine scales.

We found limited evidence of a wildfire effect on microhabitat composition immediately at (0 m distance class) and 3 m away from amphibian breeding ponds in WLNP. Although there was substantial variation among quadrats, mean values along the NMDS axes for quadrats at these distance classes did not change over time in either burned or unburned parts of the park. Likewise, although there were shifts in microhabitat composition at 0 and 3 m at most sites, there were no clear patterns in the direction of change in burned areas, and burn status was not a clear predictor of the magnitude of change in microhabitat composition for either distance class. These observations are consistent with two alternative explanations: either the processes influencing microhabitat composition immediately around amphibian breeding ponds were largely unimpacted the wildfire (i.e. high “resistance” of the system) or microhabitats were impacted by the wildfire but recovered within the three-year timeframe considered in this study (i.e. high “resilience” of the system).

The first of these explanations suggests a buffering effect of ponds on microhabitat composition during wildfire—specifically with respect to the vegetative components considered. That wetlands may play an important role in buffering terrestrial habitats during wildfire is increasingly recognized. For instance, in a remote sensing study, Fairfax & Whittle (2020) found that beaver-dammed riparian corridors remained vegetated during wildfire. However, vegetation may also recover quickly along moist pond edges, contributing to high resilience of microhabitats along pond margins. For instance, Aspinall et al. (2025) reported faster accumulation of herbaceous vegetation and development of canopy cover at a moist site than at a comparable but dry site within the Kenow burn footprint. Regardless of whether high resistance or resilience explain the current results, our results suggest that pond edges may serve as important refugia in landscapes recovering from fire, potentially enabling the nucleated recovery of wildlife populations using these refugia (e.g. Banks et al., 2017).

At the same time, our results suggest that any protection afforded to individuals by microhabitats around pond edges in fire-impacted landscapes may be limited in scale. By 10 m away from pond edges, we observed significant differences between pre-fire and post-fire microhabitat composition in the burn zone that were not observed in unburned parts of the park. Quadrats sampled 10 m away from pond edges in the burn zone following the wildfire tended to have more forbs and less moss than quadrats sampled at the same distance away from pond edges prior to the fire and, overall, most sites shifted towards a greater proportion of bare ground (represented as soil, rocks, and gravel) at this distance class. Furthermore, in a resurvey of plant communities across the park, Lloren & McCune (2024) reported greater shifts in community composition in burned parts of the park than in unburned parts of the park, consistent with a general effect of the wildfire on plant communities. Thus, away from pond margins in WLNP, microhabitats were likely much more altered.

### Notable changes at high-elevation sites

Changes in microhabitat composition around amphibian breeding ponds in the burn zone were greatest at the two highest elevation sites (1804 and 1973 m respectively) in our survey. The limited number of sites at high elevations, and specifically, the lack of sites at comparably high elevations in unburned areas, precludes clear conclusions as to the joint effects of elevation and wildfire on microhabitat composition around breeding ponds. With respect to the post-Kenow regeneration of trees, Musk et al. (2024) found that only 56% of plots in the burn zone that had trees in 1990 had seedlings in 2019/2020 after the fire (compared with 85% for unburned plots) and that the probability of seedling occurrence decreased with elevation in the burn zone. Collectively these results suggest that microhabitats around and away from high-elevation ponds may have been more heavily impacted by the Kenow wildfire than microhabitats at lower elevations and/or that recovery may be slower at higher elevations.

### Implications for amphibian populations and management

With their need for moist conditions, access to invertebrate prey, and protective cover, amphibians are highly sensitive to microhabitat conditions. Individual amphibian movements are often on the order of a few meters per day (e.g. Hoza & Bakker, 2023). Thus the fine-scale gradients in microhabitat composition away from pond edges observed here are relevant to the decisions individuals are making as they move away from breeding ponds, and may influence the speed and direction in which individuals move (e.g. Lee-Yaw et al., 2015), and ultimately the dispersion of individuals around breeding ponds during the terrestrial season (e.g. Blomquist & Hunter, 2009; Browne et al., 2009; Browne & Paszkowski, 2014; Rittenhouse & Semlitsch, 2007).

Although there were changes in microhabitat composition at and 3 m away from pond edges in burned sites, similar changes were observed at unburned sites, suggesting that the processes influencing microhabitat composition closest to pond edges were not greatly altered by the wildfire. In contrast, we detected an effect of the fire 10 m away from pond edges. One prediction that arises from the greater stability of microhabitats along pond edges relative to those further away is that, following wildfire, we may see greater effects on those species or life stages that move farther away from breeding ponds. These effects may be positive or negative depending on the population process in question. For example, western toads will move large distances away from breeding ponds provided they can find substrates with sufficient moisture and wildfires have been shown to increase western toad movement and colonization (reviewed by Hossack & Pilliod, 2011), potentially because warmer soil and burrow conditions attract toads (Hossack et al., 2009). In contrast, the loss of forest duff and moss may negatively impact long-toed salamanders, which were associated with thick litter in upslope habitats in a study farther north in Alberta (Graham, 1997) and which showed preference for moss substrates in an experimental context (Lee-Yaw et al., 2015). Even here though, it’s possible that the reduced complexity of a landscape with a greater proportion of bare ground following the fire will facilitate and encourage faster movement through the environment, at least over short distances (Lee-Yaw et al., 2015), potentially increasing the distances that individuals travel. An alternative possibility is that changes to microhabitats away from pond edges following wildfire will encourage a greater number of individuals to remain near pond edges during the terrestrial season. An increased concentration of individuals in and around wetlands may in turn may influence the frequency of both intraspecific (e.g. Kerby & Kats, 1998) and interspecific interactions (e.g. Hossack et al., 2013). Thus, differences in the amount of change in microhabitat composition observed at versus further away from pond margins in the present study leads to several testable hypotheses that would enrich our understanding of the consequences of wildfire for amphibian populations.

The changes to microhabitats observed also have implications for managemet activities in the park. For example, leopard frogs are known to avoid wooded areas (Blomquist & Hunter, 2009) and it is possible forest encroachment and densification in WLNP played a role in the extirpation of the species from the park. WLNP has been actively reintroducing the species to the park and greater forb and equisetum cover away from wetland margins may facilitate the establishment and spread of this species. In contrast, the observed changes to microhabitats at high elevation sites may be cause for concern for long-toed salamanders. Once the top-predator in these environments, the long-toed salamander has been extirpated from much of its high-elevation range in Alberta and the Rocky Mountains (Pearson, 2004). The few remaining high-elevation populations are expected to be isolated (Giordano et al., 2007) and may already exist at small population sizes (Funk et al., 1999). Pronounced changes to microhabitats at these sites may compound the risks to these populations (e.g. Hossack et al., 2013), underscoring the need for post-fire assessment and monitoring of these populations.

### Limitations and Conclusions

Our study is among the first to examine the impacts of severe wildfire on fine-scale gradients in microhabitat composition using a before-after-control-impact design. Nevertheless, as the wildfire occurred unexpectedly, the surveys were not explicitly designed to test a wildfire effect, and several limitations inherent to this retrospective analysis should be acknowledged. First, the limited number of sites originally surveyed restricted our ability to evaluate additional factors that likely contribute to variation in microhabitats across space and that could modulate the effects of wildfire (e.g. slope and/or aspect, moisture gradients, etc.). Second, we do not have concordant data on microclimatic conditions for the sites included in the current study. Although microhabitat structure is known to influence microclimates in postfire landscapes (Crockett & Hurteau, 2022; Wolf et al., 2021), the extent to which structural features of the microhabitat serve as a proxy for suitable microclimatic conditions may be limited (e.g. Evans et al., 2020). Finally, the potential implications of our findings for amphibians as discussed above, remain hypotheses that require testing. Although a parallel study detected no effect of the wildfire on site occupancy for four species in the park (Hunter, 2022), we currently lack data on processes that are closely linked to microhabitats, including individual movement and habitat selection in the park, as well as gradients in abundance. Despite these limitations, our study provides insight into how different environmental gradients can interact with wildfire to shape structural features of microhabitats. Our study further establishes a foundation for continued investigation of postfire habitat dynamics within the park and the responses, not only of amphibians, but other small-bodied species that depend on these microhabitats for shelter, foraging, and movement.

## Author Contributions

JAL conceived of the study, acquired the funding, and took the prefire photographs. TS took the postfire photographs, developed the approach to scoring microhabitat features, and collected the data. All authors developed the analytical framework, with the final analyses being completed by JAL. JAL led the writing of the manuscript with contributions from all authors.

## Acknowledgements

We thank S DeLisle, M Robinson, M Cooling, and D. Hunter for help in the field. B Johnson and K Pearson facilitated permits and access to sites in the park in 2009 and 2020 respectively.

