## Supplementary Material for "Changes in microhabitat structure around amphibian breeding ponds in the northern Rocky Mountains following severe wildfire"

**Appendix A. Supplementary material**

**Table A1.** Site information.

| Site* | # Quadrats | Burn Status | Elevation (m) | Ecoregion | Species observed** |
|---|---|---|---|---|---|
| 113 (A58) | 18 | Unburned | 1273 | Foothills Parkland | LTSA, BCFR, CSFR, WETO |
| 114 (A19) | 24 | Unburned | 1284 | Foothills Parkland | LTSA, BCFR, CSFR, WETO |
| 118 (A53) | 24 | Unburned | 1307 | Foothills Parkland | LTSA, BCFR, CSFR, WETO |
| 115 (A57) | 24 | Unburned | 1315 | Montane | LTSA, BCFR, WETO |
| 107 (A16) | 24 | Unburned | 1375 | Montane | LTSA, BCFR, CSFR, WETO |
| 108 (A50) | 24 | Unburned | 1377 | Montane | LTSA, BCFR, CSFR, WETO |
| 109 (A26) | 22 | Unburned | 1378 | Montane | LTSA, BCFR, CSFR, WETO |
| 103 (A24) | 24 | Unburned | 1527 | Montane | LTSA, BCFR, CSFR, WETO |
| 111 (A59) | 24 | Unburned | 1549 | Montane | LTSA, BCFR, WETO |
| 116 | 24 | Unburned | 1552 | Montane | - |
| 112 (A23) | 23 | Unburned | 1558 | Montane | LTSA, BCFR, CSFR, WETO |
| 117 (A3) | 24 | Burned | 1281 | Foothills Parkland | LTSA, BCFR, CSFR |
| 96 (A13) | 23 | Burned | 1295 | Foothills Parkland | LTSA, TISA, BCFR, WETO |
| 99 (A20) | 18 | Burned | 1315 | Montane | LTSA, BCFR, CSFR, WETO |
| 98 (A8) | 24 | Burned | 1323 | Montane | LTSA, TISA, BCFR, CSFR, WETO |
| 97 | 24 | Burned | 1324 | Montane | - |
| 102 (A10) | 24 | Burned | 1408 | Montane | LTSA, BCFR, WETO |
| 106 (A11) | 22 | Burned | 1672 | Lower Subalpine | LTSA |
| 105 | 24 | Burned | 1804 | Lower Subalpine | - |
| 104 (A12) | 24 | Burned | 1973 | Upper Subalpine | LTSA, CSFR |

* Names in parentheses correspond to site names used by WLNP.
** Based on multiple records in the long-term amphibian monitoring database maintained by WLNP; LTSA = long-toed salamander (*Ambystoma macrodactylum*); TISG = wester tiger salamander (*Ambystoma mavortium*); WETO = western toad (*Anaxyrus boreas*); BCFR = boreal chorus frog (*Pseudacris maculata*); CSPF = Columbia spotted frogs (*Rana luteiventris*); rows with missing values are sites where long-toed salamanders breed but where other members of the amphibian community have not been formally surveyed.

**Table A2.** Results from a PERMANOVA testing differences between microhabitat composition between 1 $m^2$ quadrats at three different distances (0, 3, 10 m) from pond edges at 20 amphibian breeding sites in southwestern Alberta prior to the 2017 Kenow wildfire.

| | **df** | **SS** | **MS** | **Pseudo-F** | **p-value*** |
|---|---|---|---|---|---|
| Distance Class | 2 | 24357 | 12179 | 2.9441 | **0.006** |
| Site (distance) | 57 | 237390 | 4164.7 | 3.0843 | **0.0002** |
| Residual | 171 | 230900 | 1350.3 | | |
| Total | 230 | 492850 | | | |

*based on 5000 permutations

**Table A3.** Average dispersion around the centroid for different distance class groups (0, 3, 10 m) in multivariate space describing microhabitat composition and results from a permutation test* of differences in dispersion among groups prior to the 2017 Kenow wildfire.

| | **df** | **F** | **p-value*** |
|---|---|---|---|
| Distance Class | 2 | 2.9985 | **0.09** |
| *Average dispersion* | | | |
| *0 m = 44.743* | | | |
| *3 m = 44.556* | | | |
| *10 m = 40.359* | | | |
| Residual | 228 | | |

*based on 5000 permutations

**Table A4.** Results from post-hoc tests assessing pairwise differences in relative microhabitat compositions among quadrats at different distances away from amphibian breeding pond edges in southwestern Alberta.

| **Distance Class 1** | **Distance Class 2** | **Average Similarity between groups** | **t** | **p-value*** |
|---|---|---|---|---|
| 0 | 3 | 37.18 | 1.3967 | 0.1156 |
| 0 | 10 | 36.453 | 2.3658 | **0.0008** |
| 3 | 10 | 40.367 | 0.99196 | 0.4231 |

*based on 5000 permutations

**Table A5.** Results from an indicator analysis examining microhabitat features significantly associated with different distances away from amphibian breeding pond edges in southwestern Alberta prior to the 2017 Kenow wildfire. Features that are significantly associated with any distance class are bolded.

| Microhabitat Feature | 0m | 3 m | 10 m | Association coefficient | p-value |
|---|---|---|---|---|---|
| Grass | 0 | 1 | 1 | 0.0814 | 0.2467 |
| **Sedge** | **1** | **0** | **0** | **0.2836** | **0.00020** |
| Forb | 0 | 1 | 1 | 0.1727 | 0.0080 |
| Equisetum | 0 | 0 | 1 | 0.06437 | 0.5547 |
| **Shrub** | **0** | **1** | **1** | **0.1916** | **0.00240** |
| **Deciduous tree** | **0** | **0** | **1** | **0.1999** | **0.00480** |
| Coniferous tree | 0 | 0 | 1 | 0.08183 | 0.3935 |
| Coarse woody debris | 0 | 1 | 1 | 0.1137 | 0.1490 |
| Snag | 0 | 1 | 0 | 0.1070 | 0.2200 |
| Rock | 1 | 0 | 0 | 0.08845 | 0.3793 |
| Moss/lichen | 1 | 0 | 0 | 0.06527 | 0.4995 |
| Litter | 1 | 0 | 0 | 0.07539 | 0.3701 |
| **Soil** | **1** | **1** | **0** | **0.1883** | **0.00100** |
| Gravel | 1 | 0 | 0 | 0.03857 | 0.7886 |

**Table A6.** Results from post-hoc tests assessing pairwise differences in relative microhabitat composition before and after a major wildfire in southwestern Alberta, in burned and unburned areas based on quadrats at different distances away from amphibian breeding ponds.

| Comparison | Average similarity between prefire and postfire samples | t | p-value* |
|---|---|---|---|
| *0 m distance class* | | | |
| Burned | 53.482 | 1.1184 | 0.307 |
| Unburned | 52.382 | 1.6859 | 0.070 |
| *3 m distance class* | | | |
| Burned | 55.493 | 1.3095 | 0.193 |
| Unburned | 57.467 | 1.0017 | 0.419 |
| *10 m distance class* | | | |
| Burned | 54.687 | 1.9826 | **0.026** |
| Unburned | 63.73 | 1.2589 | 0.199 |

*based on 5000 permutations

**Table A7.** Microhabitat features as fitted vectors onto a NMDS ordination of 1 $m^2$ quadrats (N = 462) from 20 amphibian breeding ponds in southwestern Alberta collected at three distance classes (0, 3, 10 m), two time points (2009: prefire; 2020: postfire), and from areas that were burned and unburned by the 2017 Kenow wildfire. Columns show the direction of each vector along each NMDS axis, overall strength of association ($r^2$) of each vector, and the significance of the association based on 5000 permutations. Significant vectors ($p < 0.01$) are bolded.

| Microhabitat Feature | NMDS1 | NMDS2 | NMDS3 | $r^2$ | p-value |
|---|---|---|---|---|---|
| **grass** | **-0.81418** | **0.53821** | **-0.21781** | **0.9122** | **0.0002** |
| **sedge** | **-0.16263** | **-0.4186** | **0.89349** | **0.8706** | **0.0002** |
| **forb** | **0.13373** | **-0.35102** | **-0.92677** | **0.4816** | **0.0002** |
| **equisetum** | **0.19929** | **0.34591** | **-0.91686** | **0.0695** | **0.0004** |
| **shrub** | **0.81563** | **0.5737** | **0.07497** | **0.9061** | **0.0002** |
| **deciduous tree** | **-0.04177** | **0.47108** | **-0.8811** | **0.0374** | **0.0004** |
| coniferous tree | 0.92812 | -0.08476 | -0.36251 | 0.0607 | 0.0270 |
| coarse woody debris | 0.30232 | -0.57557 | -0.75982 | 0.0461 | 0.4981 |
| snag | 0.77826 | -0.42194 | -0.46506 | 0.0645 | 0.0330 |
| **rock** | **0.026** | **-0.74996** | **-0.66097** | **0.1366** | **0.0002** |
| **moss/ lichen** | **-0.19498** | **-0.53944** | **-0.81914** | **0.0403** | **0.0064** |
| litter | -0.86868 | -0.0636 | 0.49127 | 0.0042 | 0.8136 |
| **soil** | **-0.00179** | **-0.86561** | **-0.50072** | **0.6002** | **0.0002** |
| **gravel** | **0.13733** | **-0.81526** | **-0.56258** | **0.1965** | **0.0002** |

*Permutations were restricted to quadrats within sites to preserve the structure of the data.

**Table A8.** Microhabitat features as fitted vectors onto NMDS ordinations of pre- and post-fire samples from 20 amphibian breeding ponds at different distances to aid with interpretation of Figure 4 from the main text. Columns show the direction of each vector along each NMDS axis, overall strength of association ($r^2$) of each vector, and the significance of the association based on 5000 permutations*. Significant vectors ($p < 0.01$) are bolded.

| Distance class | Microhabitat Feature | NMDS1 | NMDS2 | NMDS3 | $r^2$ | p-value |
|---|---|---|---|---|---|---|
| ***0 m*** | **grass** | **0.92893** | **0.37026** | | **0.2489** | **0.0062** |
| | **sedge** | **-0.93591** | **-0.35223** | | **0.8616** | **0.0002** |
| | forb | 0.91826 | 0.39597 | | 0.3298 | 0.0230 |
| | equisetum | 0.82418 | -0.56633 | | 0.2152 | 0.0802 |
| | **shrub** | **0.64879** | **-0.76097** | | **0.6924** | **0.0004** |
| | deciduous tree | 0.59913 | 0.80065 | | 0.0702 | 0.2569 |
| | coniferous tree | 0.99321 | 0.11637 | | 0.0699 | 0.1256 |
| | coarse woody debris | 0.86714 | 0.49807 | | 0.1705 | 0.0878 |
| | snag | 0.99813 | -0.06109 | | 0.1194 | 0.5009 |
| | rock | 0.16593 | 0.98614 | | 0.3372 | 0.1798 |
| | moss/ lichen | 0.85841 | 0.51296 | | 0.1084 | 0.6369 |
| | litter | 0.91879 | -0.39474 | | 0.083 | 0.3433 |
| | soil | -0.20989 | 0.97772 | | 0.7759 | 0.0104 |
| | gravel | -0.05229 | 0.99863 | | 0.3994 | 0.0150 |
| ***3 m*** | grass | -0.80431 | -0.5942 | | 0.2488 | 0.2038 |
| | **sedge** | **-0.56869** | **-0.82255** | | **0.7452** | **0.0008** |
| | **forb** | **-0.14056** | **0.99007** | | **0.2899** | **0.0010** |
| | equisetum | 0.47092 | -0.88217 | | 0.0698 | 0.1590 |
| | **shrub** | **0.98151** | **-0.1914** | | **0.9156** | **0.0038** |
| | deciduous tree | 0.10792 | -0.99416 | | 0.0105 | 0.8040 |
| | **coniferous tree** | **0.95604** | **0.29324** | | **0.2517** | **0.0044** |
| | coarse woody debris | 0.42669 | 0.9044 | | 0.1 | 0.5075 |
| | snag | 0.68529 | 0.72827 | | 0.2695 | 0.1312 |
| | rock | -0.43017 | 0.90275 | | 0.3814 | 0.3815 |
| | moss/ lichen | 0.34588 | 0.93828 | | 0.1414 | 0.8966 |
| | litter | 0.47471 | 0.88014 | | 0.0896 | 0.1738 |
| | **soil** | **-0.49797** | **0.8672** | | **0.6996** | **0.0002** |
| | gravel | -0.42338 | 0.90595 | | 0.4995 | 0.3171 |
| ***10 m*** | **grass** | **-0.54305** | **0.30814** | **-0.78112** | **0.8099** | **0.0006** |
| | **sedge** | **0.5475** | **-0.46874** | **-0.6932** | **0.7239** | **0.0006** |
| | **forb** | **-0.24965** | **-0.7641** | **0.59483** | **0.4754** | **0.0018** |
| | equisetum | 0.65398 | 0.71032 | 0.2603 | 0.1708 | 0.0280 |
| | **shrub** | **0.61168** | **0.61185** | **0.50148** | **0.8969** | **0.0002** |
| | deciduous tree | 0.3708 | 0.01436 | -0.9286 | 0.3147 | 0.6711 |
| | coniferous tree | -0.78229 | 0.46202 | 0.4178 | 0.446 | 0.0088 |
| | coarse woody debris | -0.9229 | -0.17912 | 0.34084 | 0.4687 | 0.0152 |
| | snag | -0.29672 | -0.72747 | 0.61866 | 0.1973 | 0.1902 |
| | **rock** | **-0.08394** | **-0.97993** | **0.18079** | **0.5053** | **0.0080** |
| | moss/ lichen | 0.25097 | -0.45079 | 0.85662 | 0.1409 | 0.7728 |
| | litter | 0.13413 | 0.23664 | 0.96229 | 0.0833 | 0.8480 |
| | **soil** | **0.08786** | **-0.9446** | **0.31625** | **0.6631** | **0.0054** |
| | gravel | -0.27995 | -0.75482 | 0.5932 | 0.1742 | 0.3805 |

*Permutations were restricted to samples within sites to preserve the structure of the data.

**Table A9.** Sensitivity of linear models (beta regression) assessing predictors of change in microhabitat composition (pairwise Bray-Curtis dissimilarity) around amphibian breeding ponds in southwestern Alberta to the exclusion of high-elevation sites in the burn zone that had high leverage in the models[*].

| | | Without site 104 | | | | Without sites 104 and 105 | | | | Without sites 104, 105, and 106 | | | |
|---|---|---|---|---|---|---|---|---|---|---|---|---|---|
| **Response** | **Predictor** | **Coef[a]** | **SE** | **Z** | **p[b]** | **Coef[a]** | **SE** | **Z** | **p[b]** | **Coef[a]** | **SE** | **Z** | **p[b]** |
| Change 0 m away from pond edges | Intercept | -0.871 | 0.192 | -4.543 | | -1.0178 | 0.202 | -5.034 | | -0.831 | 0.350 | -2.374 | |
| | Burn status | 0.0599 | 0.249 | 0.241 | 0.273 | 0.229 | 0.252 | 0.909 | 0.572 | 0.0655 | 0.384 | 0.171 | 0.503 |
| | Elevation | 0.00158 | 0.000978 | 1.614 | 0.22 | -0.000281 | 0.00157 | -0.179 | 0.630 | 0.00269 | 0.00520 | 0.517 | 0.568 |
| | Burn status x Elevation | -0.00287 | 0.00176 | -1.631 | 0.109 | -0.00102 | 0.00210 | -0.483 | 0.625 | -0.00398 | 0.00539 | -0.739 | 0.449 |
| | *Pseudo-R2* | *0.1499* | | | | *0.08692* | | | | *0.09233* | | | |
| Change 3 m away from pond edges | Intercept | -0.572 | 0.129 | -4.42 | | -0.684 | 0.136 | -5.04 | | -0.476 | 0.233 | -2.045 | |
| | Burn status | -0.376 | 0.174 | -2.166 | **0.0109** | -0.245 | 0.176 | -1.393 | 0.191 | -0.437 | 0.259 | -1.684 | 0.246 |
| | Elevation | 0.00211 | 0.000675 | 3.128 | **0.0140** | 0.000986 | 0.00103 | 0.96 | 0.403 | 0.00478 | 0.00348 | 1.373 | 0.267 |
| | Burn status x Elevation | -0.00311 | 0.00125 | -2.483 | **0.0184** | -0.00199 | 0.00145 | -1.37 | 0.178 | -0.00579 | 0.00363 | -1.594 | 0.128 |
| | *Pseudo-R2* | *0.4476* | | | | *0.1765* | | | | *0.1964* | | | |
| Change 10 m away from pond edges | Intercept | -0.868 | 0.177 | -4.904 | | -0.980 | 0.190 | -5.147 | | -0.822 | 0.333 | -2.472 | |
| | Burn status | 0.0353 | 0.229 | 0.154 | 0.239 | 0.160 | 0.239 | 0.669 | 0.562 | 0.0134 | 0.36532 | 0.037 | 0.489 |
| | Elevation | 0.00217 | 0.000894 | 2.422 | 0.0781 | 0.00100 | 0.00142 | 0.704 | 0.706 | 0.00393 | 0.00497 | 0.791 | 0.672 |
| | Burn status x Elevation | -0.00278 | 0.00161 | -1.731 | 0.091 | -0.00162 | 0.00195 | -0.832 | 0.408 | -0.00455 | 0.00515 | -0.883 | 0.393 |
| | *Pseudo-R2* | *0.2236* | | | | *0.06568* | | | | *0.08239* | | | |

[*] Each iteration of the model revealed a single high-leverage site, starting with the original model which identified site 104 as such; results here show effects of sequential removal of the three highest elevation sites in the burn zone.

[a] Coefficients for burn status represent comparison to the reference category “unburned”.

[b] p-values reflect marginal effects of predictor and were obtained from likelihood ratio tests comparing model with and without the predictor; to test each main effects, both the main effect and the interaction term were removed in the reduced model. Significant values are bolded.

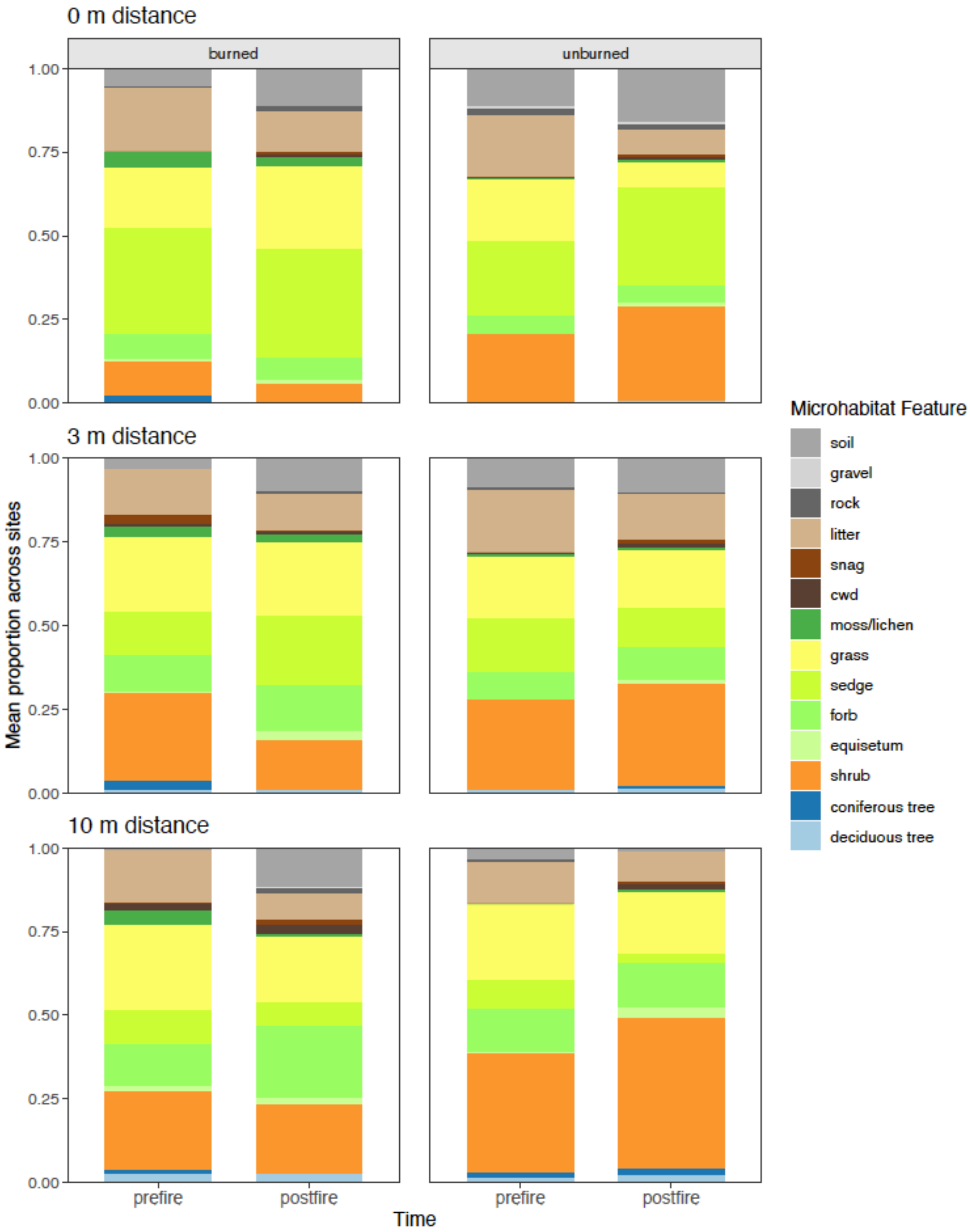


**Figure A1.** Stacked barplots showing the total proportion of counts for each microhabitat feature at three different distances away from pond edges for burned and unburned sites before and after the 2017 Kenow wildfire.

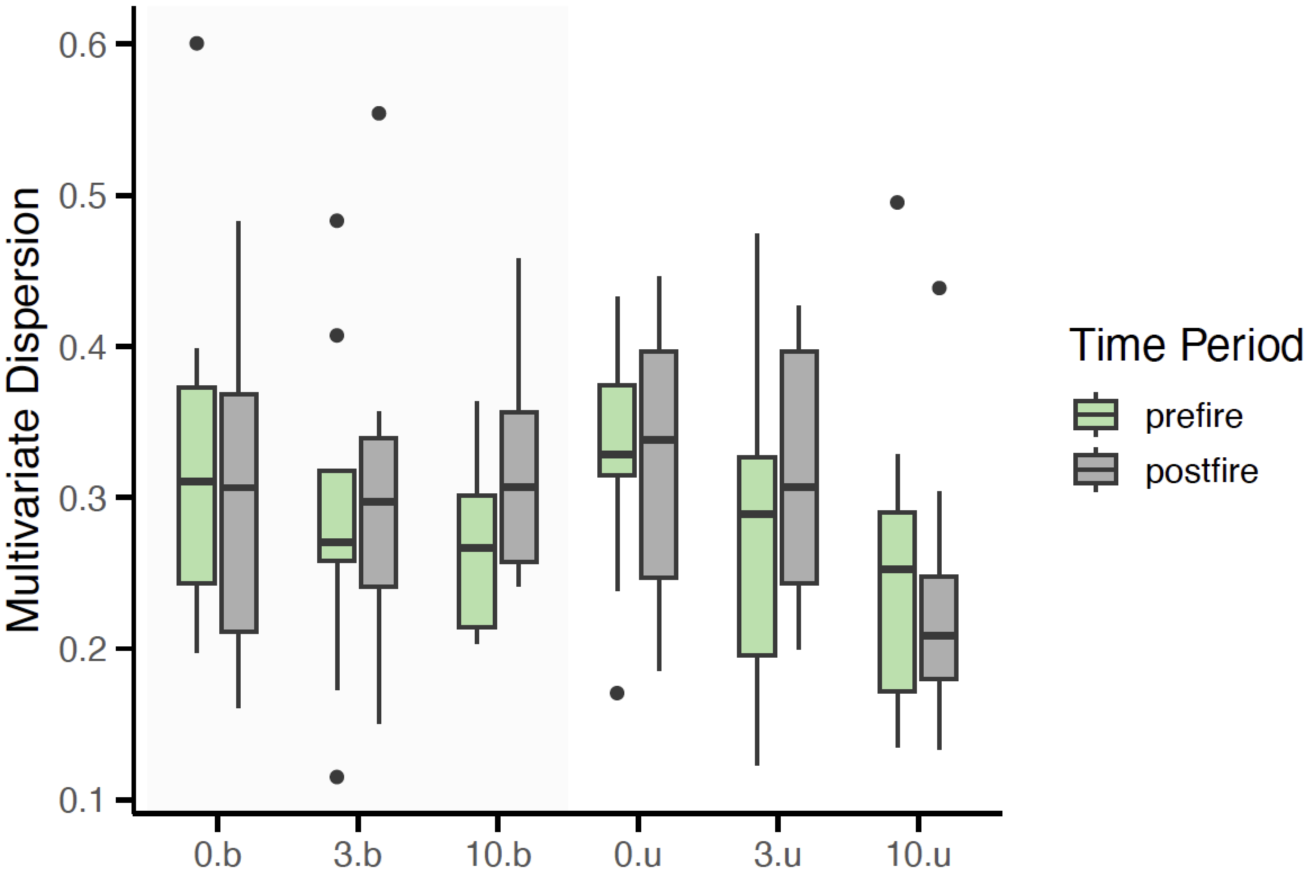


**Figure A2.** Boxplots of multivariate dispersion (distance to group centroid) for quadrats at different distances away from pond edges (0, 3, 10 m) from burned (left, shaded) and unburned (right, unshaded) sites in southwestern Alberta. Dispersion is based on pairwise Bray-Curtis dissimilarities between quadrats.
